# webMCP-counter: a web interface for transcriptomics-based quantification of immune and stromal cells in heterogeneous human or murine samples

**DOI:** 10.1101/2020.12.03.400754

**Authors:** Maxime Meylan, Etienne Becht, Catherine Sautès-Fridman, Aurélien de Reyniès, Wolf H. Fridman, Florent Petitprez

## Abstract

**Summary:** We previously reported MCP-counter and mMCP-counter, methods that allow precise estimation of the immune and stromal composition of human and murine samples from bulk transcriptomic data, but they were only distributed as R packages. Here, we report webMCP-counter, a user-friendly web interface to allow all users to use these methods, regardless of their proficiency in the R programming language.

**Availability and Implementation:** Freely available from http://134.157.229.105:3838/webMCP/. Website developed with the R package shiny. Source code available from GitHub: https://github.com/FPetitprez/webMCP-counter.

## Introduction

In many areas of biology, it is crucial to quantify the composition of heterogeneous samples at the cell type level. For instance, tumors are highly heterogeneous systems harboring not solely cancer cells, but also a wide variety of immune and stromal cells^1^. The extent and composition of this immune infiltrate has been repeatedly shown to be associated with patients’ survival^2^ and response to treatment^3^. Bulk transcriptomics capture signals from all cell types in a sample and can therefore be used to estimate its composition through deconvolution methods^4^. Several methods have been reported to estimate the immune and/or stromal composition of human samples, and only a few for murine samples^5^.

Our group developed MCP-counter^6^ and its murine counterpart mMCP-counter^7^, which were distributed as R packages. However, using these methods requires proficiency in R programming. To open the use of these methods to all biologists, we report here the development of webMCP-counter, a user-friendly web interface for both MCP-counter and mMCP-counter. This tool can be accessed using this url: http://134.157.229.105:3838/webMCP/

## webMCP-counter

Our web interface, webMCP-counter, is composed of two main steps (Figure 1). The first step aims at running (m)MCP-counter per se. This step requires the user to upload a transcriptomics dataset that can be presented under various formats. The user also needs to specify several details regarding the dataset (file format, gene ID format and organism). Either HUGO gene symbol or ENSEMBL ID are valid formats for genes. Depending on the organism, either MCP-counter (human data) or mMCP-counter (mouse data) is run, the scores are presented as a table that can be visualized within the tool or downloaded for other purposes.

**Fig. 1.**
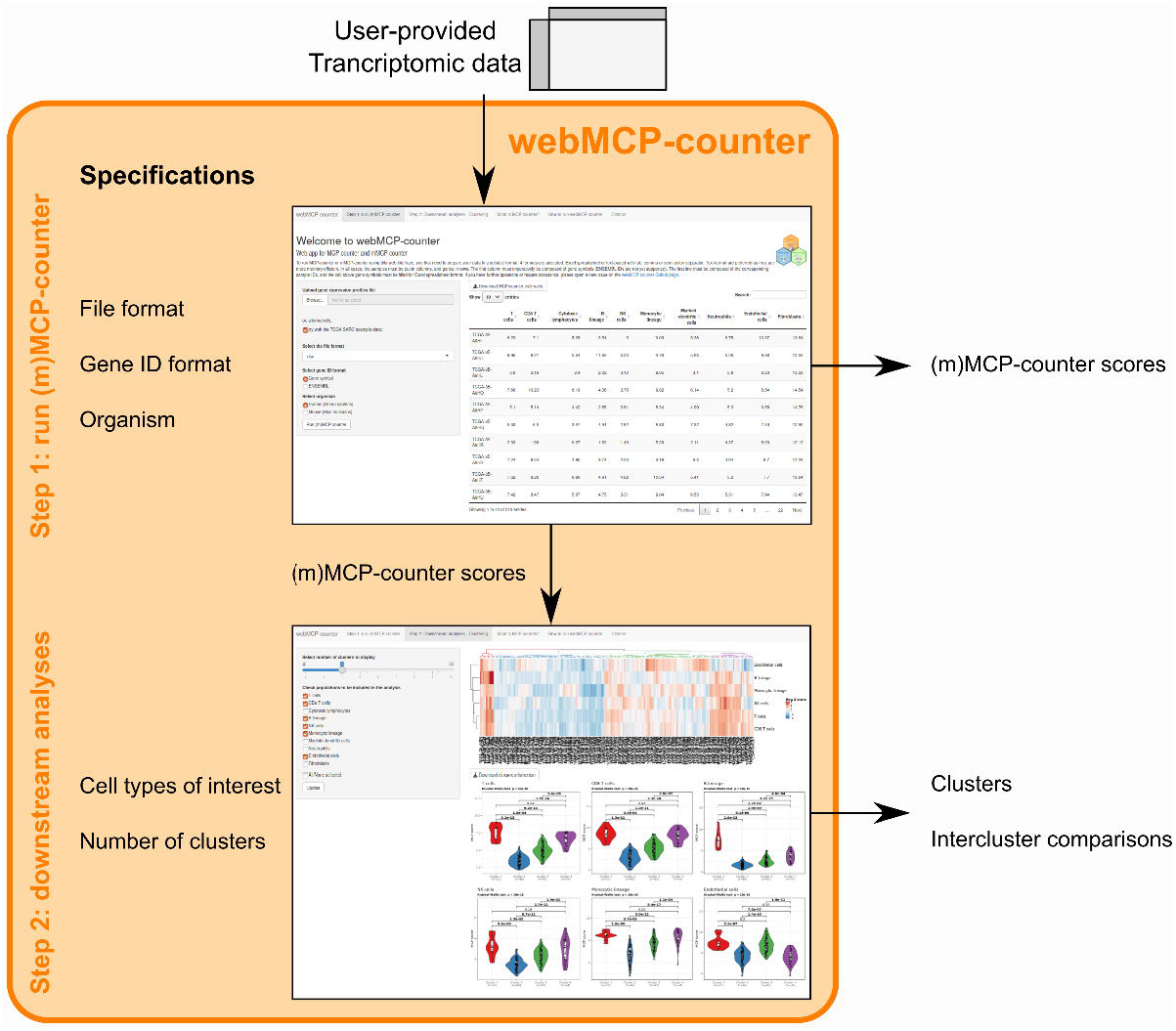
Diagram explaining the main steps of a typical analysis using webMCP-counter. In the first step, the user uploads transcriptomic data and sets the parameters. (m)MCP-counter is then run on the user-provided data and can be accessed as output and passed on to the second step. Here, a clustering is performed based on the user-selected cell populations and number of classes.

The second step consists in a typical downstream analysis. The user is prompted to choose a number of clusters and the cell populations to include in the analysis among the populations available, which depends on the organism and the data provided. Then, a hierarchical clustering with standard parameters (Euclidian distance and Ward linkage criterion) is run on the (m)MCP-counter Z-scores limited to the requested populations is run. The output is displayed as a heatmap and individual violin plots detailing the differences between clusters for each selected population, and tested with a Kruskal-Wallis test followed by Benjamini-Hochberg-corrected Dunn test for pairwise comparisons, or a Mann-Whitney test for two clusters. The assignment of samples to clusters can be downloaded as a tabular file. Our tool is, to the best of our knowledge, the only one to offer to perform easily this downstream analysis from the MCP-counter output for users who are not proficient in the R programming language.

## Example on the TCGA SARC dataset

For users who simply wish to test the possibilities offered by webMCP-counter, we integrated an example dataset from the TCGA SARC cohort^8^. This dataset is composed of n=213 tumors of soft-tissue sarcoma patients analyzed by RNA-seq. By selecting the example dataset on the webMCP-counter (Figure 2, marker 1), all other parameters are internally set to the correct values, and the user-provided parameters are ignored.

**Fig. 2.**
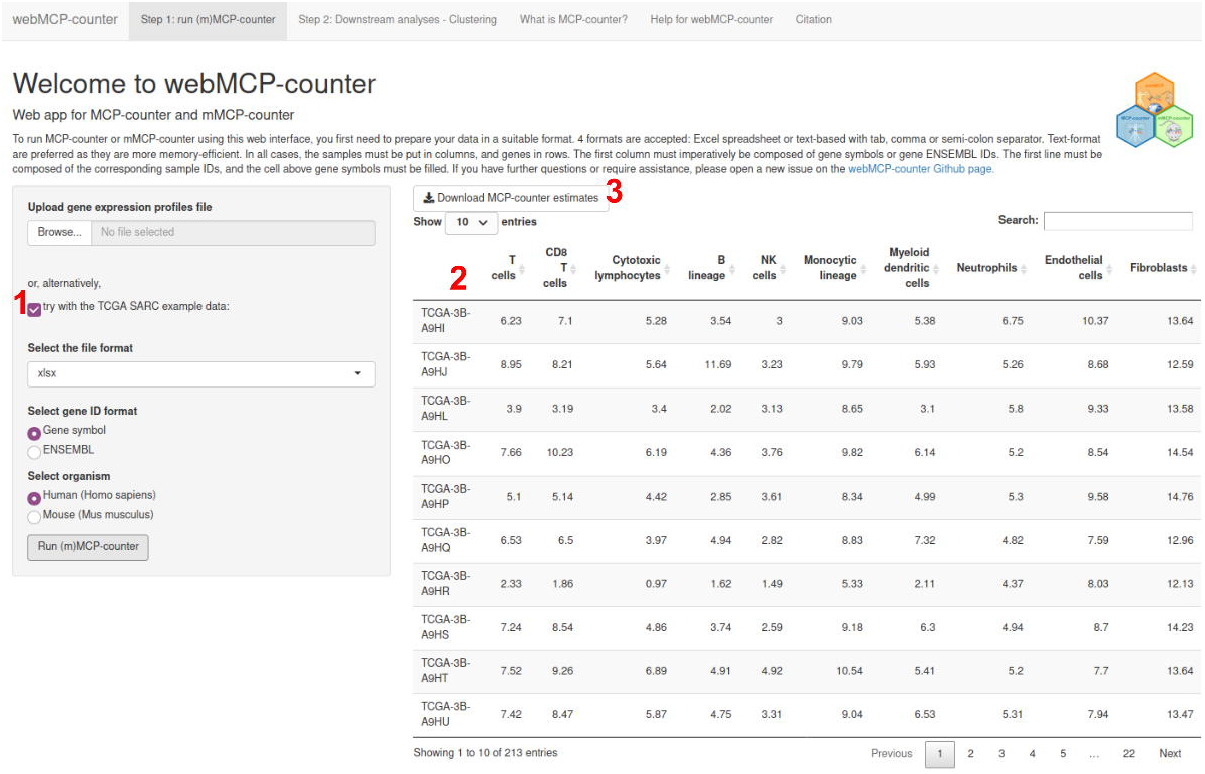
Step 1 of the analysis of the TCGA SARC example dataset. Marker 1 indicates the box to tick to access the example dataset instead of a user-provided dataset. 2 indicates the MCP-counter scores table, accessible after MCP-counter has been run. Finally, 3 indicates the download link for the .csv file containing the full table.

On the step 1 tab, users obtain a table with the MCP-counter scores for each sample in the TCGA SARC cohort (Figure 2, marker 2). This table is interactive and users can sort samples on increasing or decreasing order for each population. It is also possible to search for one particular sample by entering its sample name in the search bar. It is important to note that scores should not be interpreted as absolute proportions of cell populations in the samples, but are instead expressed in arbitrary units, similar to gene expression values. However, they can robustly be used to compare the abundances of the cell populations between different samples, such as what has been done in several studies^9–11^. This table can be downloaded as comma-separated values table (.csv) for use in other software, such as Microsoft Excel (Figure 2, marker 3).

On the Step 2 tab (Figure 3), users can perform a classical downstream analysis from the MCP-counter scores table, sample clustering based on their immune and stromal composition. To use this tab, the first step must be completed beforehand. In this tab, the user is prompted to select the populations of interest on which to perform hierarchical clustering (marker 1) and the number of clusters (marker 2). In the example of Figure 3, the clustering was performed on T cells, CD8^+^ T cells, B lineage, NK cells, monocytic lineage and endothelial cells, and 4 clusters have been required.

**Fig. 3.**
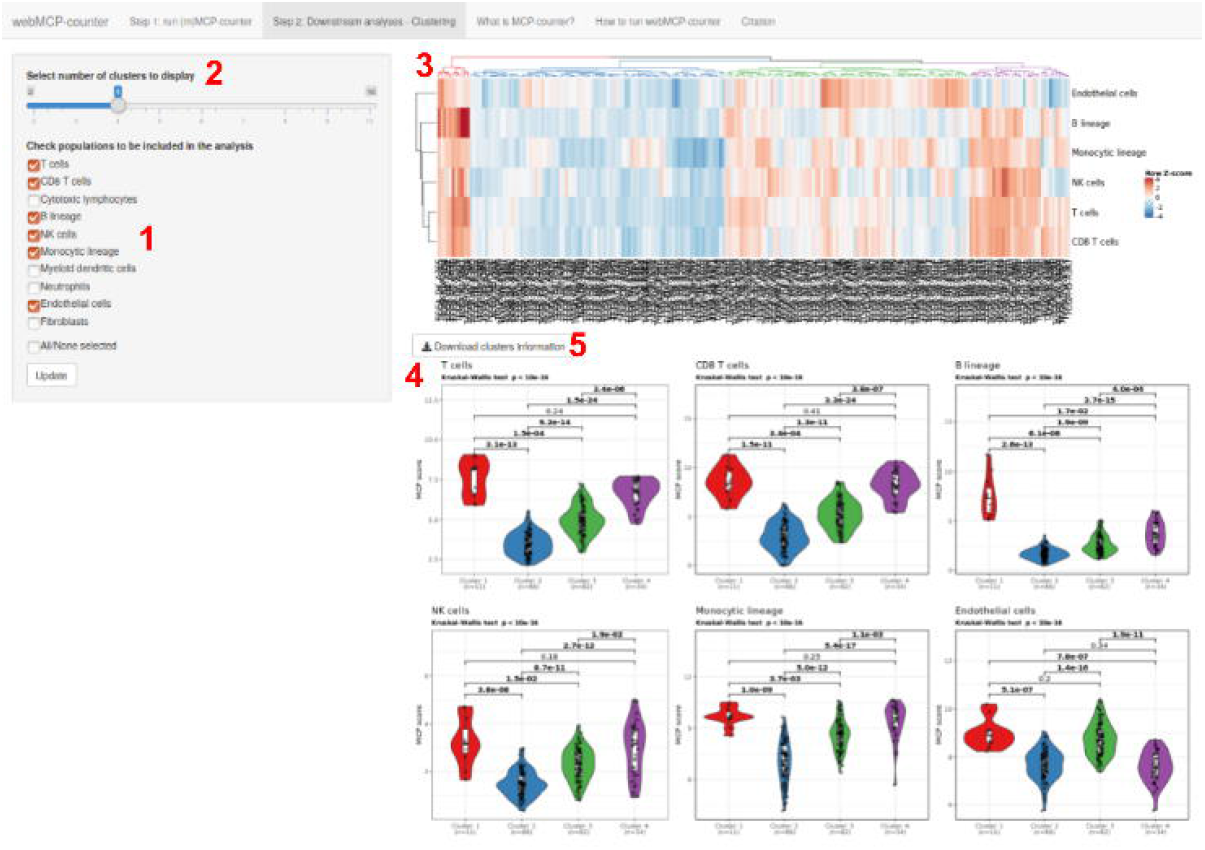
Step 2 of the analysis of the TCGA SARC example dataset. 1 marks the selection of the populations of interest. 2 indicates the selection of the number of clusters. 3 indicates the heatmap output. 4 marks the violinplots/boxplots output. 5 marks the download link for the clusters information file.

Two graphical outputs are provided when the “update” button is selected: a heatmap (marker 3) and boxplots for all selected populations (marker 4). Both can easily be downloaded via a right click in the image. The clusters information, detailing the cluster of belonging of each sample, can be downloaded by clicking on the appropriate link (marker 5).

The heatmap (Figure 3, marker 3) represents the cell population Z-scores for each samples. The clustering is computed using the Euclidian metric and Ward linkage criterion. Both samples and cell populations are clustered.

The violin plots (marker 4) output details the MCP-counter scores for all cell population of interest depending on the clusters presented in the heatmap. If one or more clusters has less than 3 samples, violin plots are replaced by boxplots. For each cell population, the relationship between the amount of this population and the clusters is estimated with a Kruskal-Wallis test. All pairwise comparisons are estimated by a post-hoc Dunn test and all p-values are indicated in the plots. P-values below the classical 0.05 threshold for significance are indicated in bold.

## Comparison with other methods

Several other methods to deconvolute the immune and stromal composition from transcriptomic data of human tissues have been previously reported. In a benchmark study published last year comparing human-based methods^12^, MCP-counter was found to be among the top-scoring methods for both accuracy (measured through correlation between the scores and randomized mixtures compositions) and specificity (measured through “spillover effect”). Murine counterparts for these methods are scarcer. In the article reporting the development of mMCP-counter, we demonstrated on an independent dataset that it outperformed pre-existing methods^7^. Importantly, our set of methods is the only one to allow a similar analysis of human and murine samples, which can prove extremely valuable for comprehensive studies that include murine models and human samples. webMCP-counter includes methods that are fast-computing, as they rely on the computation of the mean expression of gene signatures and do not require more complex data processing.

## Software implementation and availability

webMCP-counter was developed as a Shiny application in R^13^. The heatmap generation is carried using the ComplexHeatmap package^14^. Other used packages include dendextend^15^, ggplot2^16^ and dunn.test. All source code for webMCP-counter, placed under copyleft GNU license, is available from GitHub: https://github.com/FPetitprez/webMCP-counter. Users wishing to run webMCP-counter locally can download it from GitHub, install dependencies and run it as a regular shiny application.

## Conclusion

In many areas of immunology, it is crucial to accurately estimate immune and stromal composition of heterogeneous samples, such as the tumor microenvironment in cancer immunology. To this end, we previously reported MCP-counter and mMCP-counter to quantify cell populations in human and murine samples, respectively, using bulk transcriptomic data. However, these tools were only accessible to R users. By developing webMCP-counter, we offer the possibility to all users to access our methods and apply them on their datsets.

## Acknowledgements

We thank our colleagues who helped us test webMCP-counter.

